# Combating the scientific decline effect with confidence (intervals)

**DOI:** 10.1101/034074

**Authors:** David M. Groppe

**Author notes:** **Corresponding Author:** David M. Groppe Department of Psychology University of Toronto 100 St. George St. Toronto, ON M5S 3G3 Canada.

## Abstract

A symptom of the need for greater reproducibility in scientific practice is the “decline effect,” the fact that the size of many experimental effects decline with subsequent study or fail to replicate entirely. A simple way to combat this problem is for scientists to more routinely use confidence intervals (CIs) in their work. CIs provide frequentist bounds on the true size of an effect and can reveal when a statistically significant effect is possibly too small to be reliable or when a large effect might have been missed due to insufficient statistical power. CIs are often lacking in psychophysiological reports, likely due to the large number of dependent variables, which complicates deriving and visualizing CIs. In this article, I explain the value of CIs and show how to compute them for analyses involving multiple variables in various ways that adjust the intervals for the greater uncertainty induced by multiple statistical comparisons. The methods are illustrated using a basic visual oddball event-related potential (ERP) dataset and freely available Matlab software.

## Introduction

A few years ago experimental psychologist Jonathan Schooler (Schooler, 2011) published a high-profile essay on the fact that many empirical phenomena decline in size over the course of scientific studies, sometimes to the point of failing to replicate entirely. A recent attempt to replicate 100 psychology experiments suggests that this “scientific decline effect” is shockingly prevalent in experimental psychology (Open Science Collaboration, 2015). Specifically, this study found that less than 40% of the findings replicated, despite including sufficient participants to have a high likelihood of replication. Moreover, 83% of the replicated effects were smaller than initially reported. As Schooler notes, many factors potentially contribute to this phenomenon. These include:

- Regression to the mean (i.e., self-correction of an initially anomalously large outcome)
- The failure to report null results
- The failure to report observations that are inconsistent with a hypothesis
- The addition or removal of observation, variables, or analyses to generate statistical significance

To better understand and counter the decline effect, Schooler suggests pre-registering studies and publishing subsequently collected data in an open-access database. This would surely improve the reproducibility of scientific findings and since Schooler’s essay, there have been some steps towards making such databases a reality in psychophysiology. A few neuroscience and psychology journals (e.g., *Cortex, Perspectives on Psychological Science, Experimental Psychology*, and *AIMS Neuroscience*) now support the “Registered Reports” model of publishing scientific findings. In this model, a study is submitted to a journal before data are collected. If the study’s design and motivation are approved by reviewers, then the study is guaranteed publication once the data are acquired, regardless of the outcome. In addition, the Open Science Framework (https://osf.io/), provides free study pre-registration independent of journals. However, it is unlikely that this research paradigm will become mainstream unless there are greater incentives for scientists to adopt it, which is a considerable challenge (Chambers et al., 2015). Moreover, it doesn’t address the contribution of regression to the mean to the decline effect.

A more readily implementable way to combat the decline effect that does address regression to the mean is to require investigators to report confidence intervals (CIs) that will frequently span the true value of any effects. CIs are a basic tool of inferential statistics but are under-utilized in psychophysiology as researchers frequently focus solely on *p*-values. In other words, psychophysiologists often simply report that there is good evidence of a relationship between two variables (i.e., the effect has a small *p*-value) but fail to provide explicit estimates of the size of the effect.

While there is a close relationship between *p*-values and CIs (the smaller the *p*-value, the greater the distance between the confidence interval boundaries and the value of the null hypothesis being tested), deriving one from the other is often not straightforward. Thus, when only *p*-values are reported, one only clearly gets a sense of how likely there is to be some relationship between two variables, but one has little sense of how large that relationship is likely to be.

Not reporting CIs is particularly likely to lead to considerably overestimating effect sizes when studies are “underpowered.” An underpowered study is one in which the sample size is too small to be likely to detect the effects being studied and consequently the observed effect size will be highly variable (Gelman & Weakliem, 2009). When studies are underpowered and one is fortunate enough to detect an effect, the magnitude of the estimated effect is necessarily going to be much larger than the true effect since it needs to be large to qualify as statistically significant; a statistical phenomena known as the “winner’s curse” (Button et al., 2013).

To illustrate, consider a hypothetical relationship between a cup of coffee and IQ test performance. Say that in reality, a cup of coffee increases IQ test performance by 3 points on average. Since the IQ test has been designed to have a mean of 100 and a standard deviation of 15 points across the population of test takers, this amounts to a true post-coffee mean of 103 points and an effect size that is 20% of the standard deviation of the measurement noise (a small effect by Cohen’s standards—Cohen, 1988). Imagine that we perform an experiment to determine if a cup of coffee affects IQ test performance three times, each time using a different number of participants (4, 36, and 100). Imagine also that each time we do the experiment, we find that caffeine does improve test performance and get the exact same *p*-value of 0.01.

These hypothetical results are illustrated in Figure 1, which shows that when the sample is small, the effect of coffee is dramatically overestimated. If only the mean effect of coffee and the *p*-value of 0.01 had been reported in the experiment with four participants, one might interpret this as good evidence that a simple cup of coffee can increase IQ by around 19 points (enough to bump someone of average IQ up to the 90th percentile!). Indeed one might mistakenly think that the evidence from the experiment with only four participants is just as compelling as the evidence from the two larger experiments since their *p*-values are equal. However, the CIs on the bar graph help to avoid such fallacies. In two of the three scenarios, the CIs accurately span the true effect. Although the CIs still overestimate the true effect size in the experiment with four participants, the bounds nonetheless provide a reasonable sense of what the true effect of coffee might be. Moreover, one directly sees that the estimate of the size of the effect is highly imprecise when there are only four participants and one can clearly evaluate the precision of effect size estimation across the three experiments. Finally, if the experiments had turned out differently and not produced significant results, CIs would let us know if we had potentially missed an effect because our experiment had too few participants.

**Figure 1:**
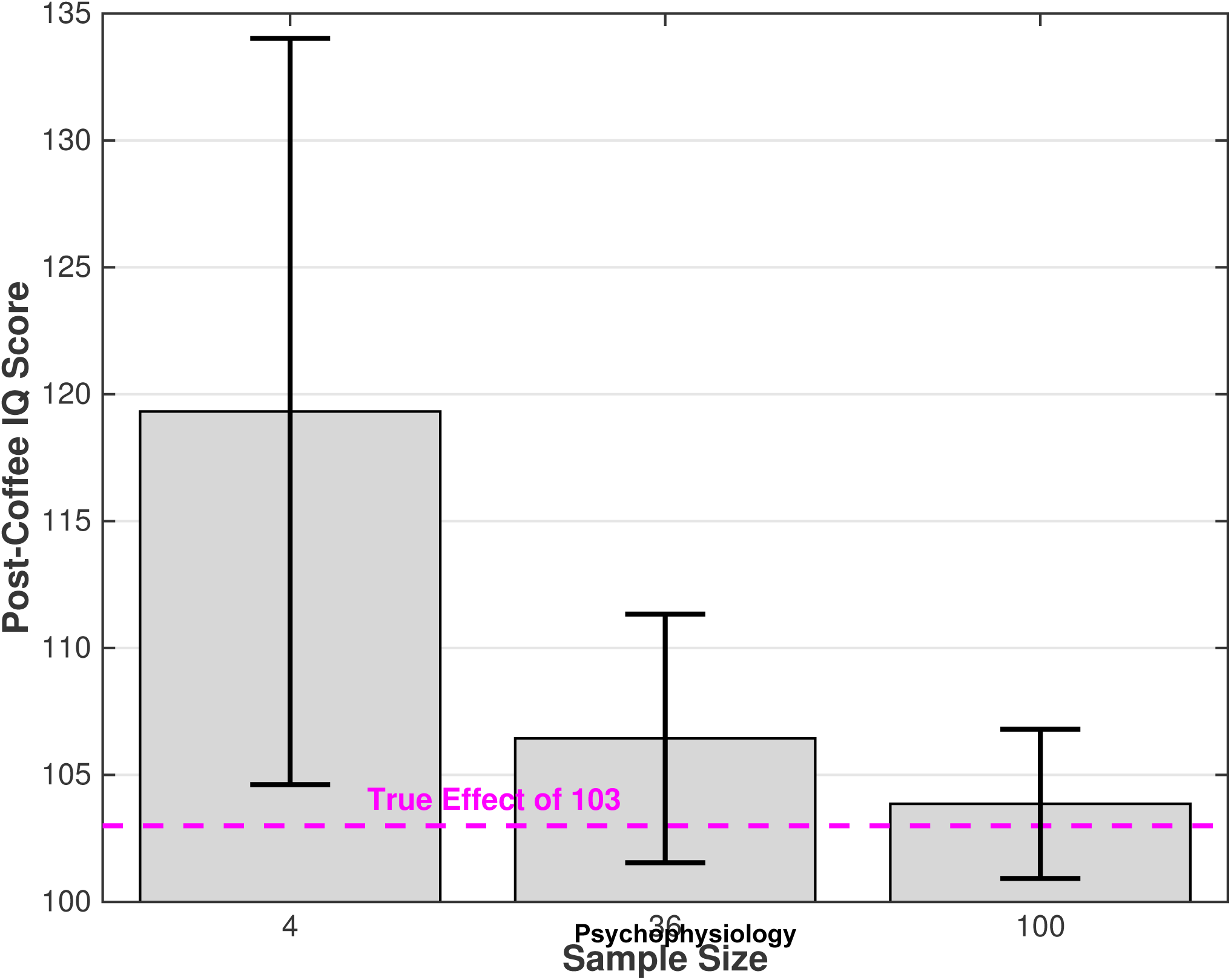
Hypothetical effect of coffee on IQ test performance in three experiments with different sample sizes. All effects have a *p*-value of 0.01. Error bars indicate 95% CIs. Magenta dashed line indicates the true mean post-coffee IQ test performance of 103.

It is important to point out that when studies are underpowered, it is unlikely that effects will be detected. Indeed, when performing the hypothetical coffee/IQ experiment with only four participants, there is only a 6.9% chance of producing a significant test result (*p*<0.05, assuming a two-tailed test). However, small effects are likely to be very prevalent in psychophysiology due to our often complicated/noisy measures of behavior and brain function and due to the large number of comparisons typical of many types of neural data (e.g., EEG, fMRI, optical imaging). Indeed, meta-analyses of the neuroscience literature suggest that the average statistical power of neuroscience experiments is at best 8 to 31% (Button et al., 2013). Thus, given the likely prevalence of small effects, the large number of studies being executed across the globe, and the fact that low power studies are particularly susceptible to bias (e.g., the addition or removal of observations and variables to generate statistical significance), the potential for overestimation of small effect sizes is surely high. Indeed, a well-known meta-analysis of fMRI studies performed by Vul and colleagues (2009) found that the scientific literature was biased to overestimate the magnitude of correlations between fMRI activation and measures of emotion, personality and social cognition.

I suspect that the reason CIs are not used more widely in psychophysiology is that historically it has been difficult to compute and represent CIs of physiological measures that often consist of thousands of dependent variables. However, with conventional data analysis and visualization software, adding CIs to analyses of physiological measures should generally be feasible. In the remainder of this article, I review a few different methods for deriving CIs in various ways that adjust the intervals for the greater uncertainty induced by multiple statistical comparisons. I illustrate the methods using a simple event-related potential (ERP) dataset. Matlab software for implementing these methods is provided as part of the freely available Mass Univariate ERP Toolbox (http://openwetware.org/wiki/Mass_Univariate_ERP_Toolbox).

## B. Computing Confidence Intervals with and without Correction for Multiple Comparisons

I will illustrate the use of CIs for ERP analysis using a simple visual oddball paradigm. These data were acquired from 16 participants at 250 Hz while they read a series of words presented one at a time on a computer monitor. The words appeared in all uppercase or lowercase letters (e.g., “ZEBRA” or “zebra”) and one type of script occurred much more frequently than the other (80% vs 20%). The type that occurred more frequently was balanced across blocks of the experiment and participants were instructed to press a button whenever they saw words in the infrequent script. I will refer to the stimuli shown in the frequent and infrequent script as “standards” and “targets,” respectively. Full experiment details have been published previously (Groppe, 2007).

Figure 2:A illustrates the grand average ERPs to targets and standards at the vertex channel. 95% CIs are illustrated with dotted lines for each time point and were derived by assuming that these ERPs are *t*-distributed. Specifically, we first obtain the *t*-scores from a *t*-distribution with 15 degrees of freedom (i.e., the number of participants minus 1) that span the central 95% of the distribution: t_0 95_(15)=±2.13. Then for each time point,z, we obtain the confidence interval by multiplying those *t*-scores by the standard error, *s*_z_, of that time point and adding it to the the mean of that time point, 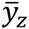:

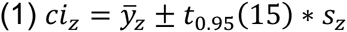

**Figure 2:**
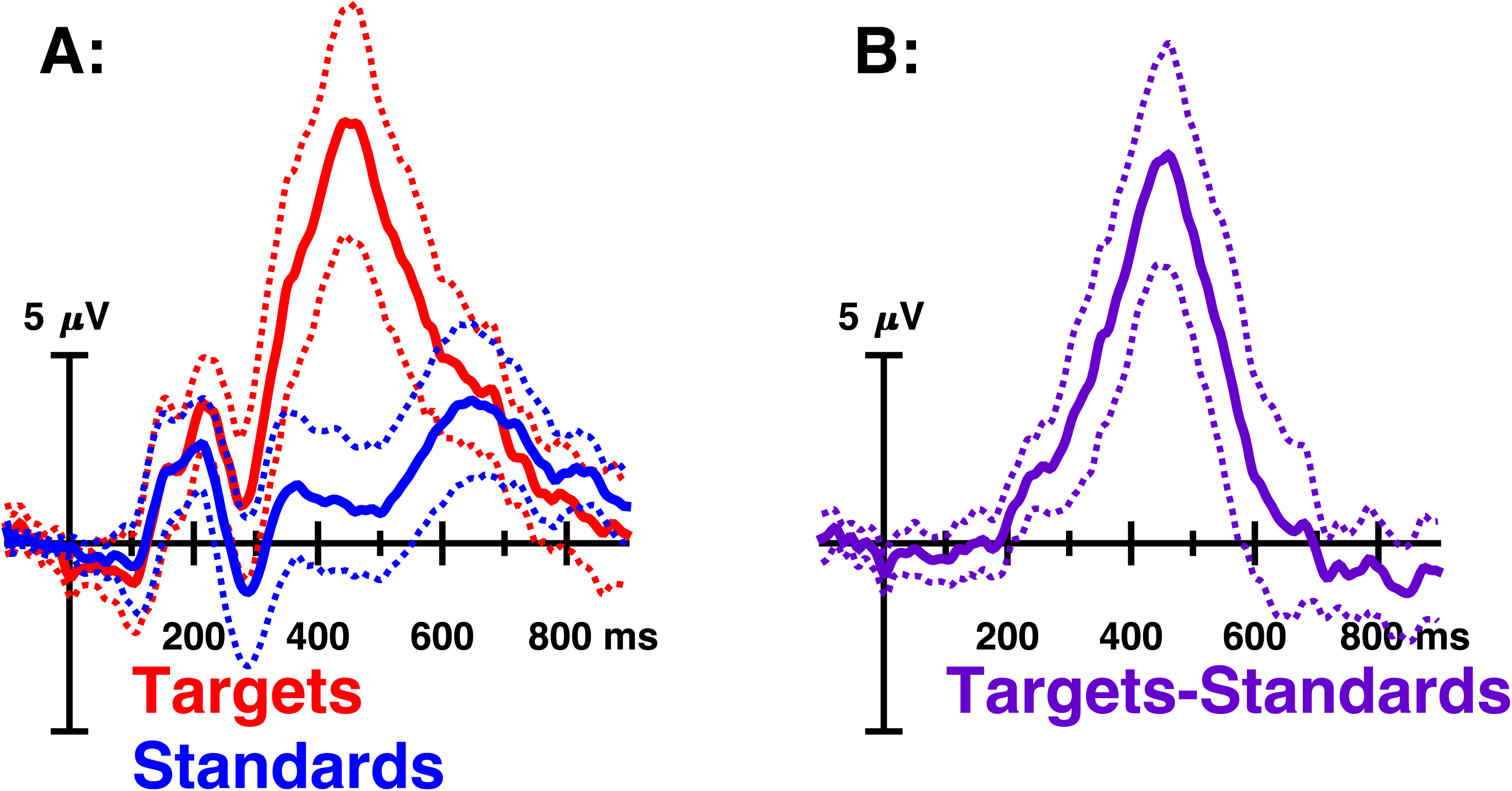
[A] Solid lines represent ERPs to an infrequent target and frequent “standard” category of text at the vertex channel. Dashed lines indicate 95% CIs with no adjustment for multiple comparisons. A large P300 effect is evident that peaks around 450 ms. [B] The difference wave and corresponding 95% CIs obtained by subtracting ERPs to standards from that to targets.

The difference between conditions is more clearly illustrated by the difference wave shown in Figure 2:B. CIs were derived as in Figure 2:A and suggest that the greater positivity to targets, a P300 effect, around 450 ms is not only a real effect (i.e., *p*<0.05) but is extremely robust. Note also that the variability across participants is not uniform across time, being greatest around the peak of the P300 effect and smallest near the baseline.

One cannot draw strong conclusions from this analysis though, because our CIs do not compensate for the large number of comparisons (in this case 351 time points) that increase our chances of finding spurious differences between conditions and consequently increase the uncertainty in our estimated differences between conditions. Several techniques such as Bonferroni correction, resampling tests, and false discovery rate (FDR) control exist for correcting for multiple comparisons when deriving *p*-values (for review see Groppe, Urbach, & Kutas, 2011a). We can use analogous techniques to adjust the size of our CIs to reflect our greater uncertainty.

### Bonferroni-Corrected Confidence Intervals

The easiest way to correct for multiple comparisons is Bonferroni correction, which simply divides our desired alpha level across the whole family of tests, *α_family_*, by the number of tests, *m*, to get the alpha level for each individual test, *α_test_*:

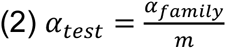

This allows us to correct our CIs by choosing more extreme *t*-scores that span 1 − *α_test_* of the *t*-distribution. For example, if we want to derive 95% CIs for every time point from 100 to 700 ms in our visual oddball experiment we are performing 151 hypotheses tests (i.e., a test for every time point between 100 and 700 ms). Thus, the *t*-scores for our CIs for each time point need to span 99.97% (i.e., 1-0.05/151) of the *t*-distribution, which would be ±4.28. We then plug those *t*-scores into Equation 1 and get the CIs illustrated in Figure 3:B, which are considerably larger than those in Figure 3:A that do not compensate for multiple comparisons.

**Figure 3:**
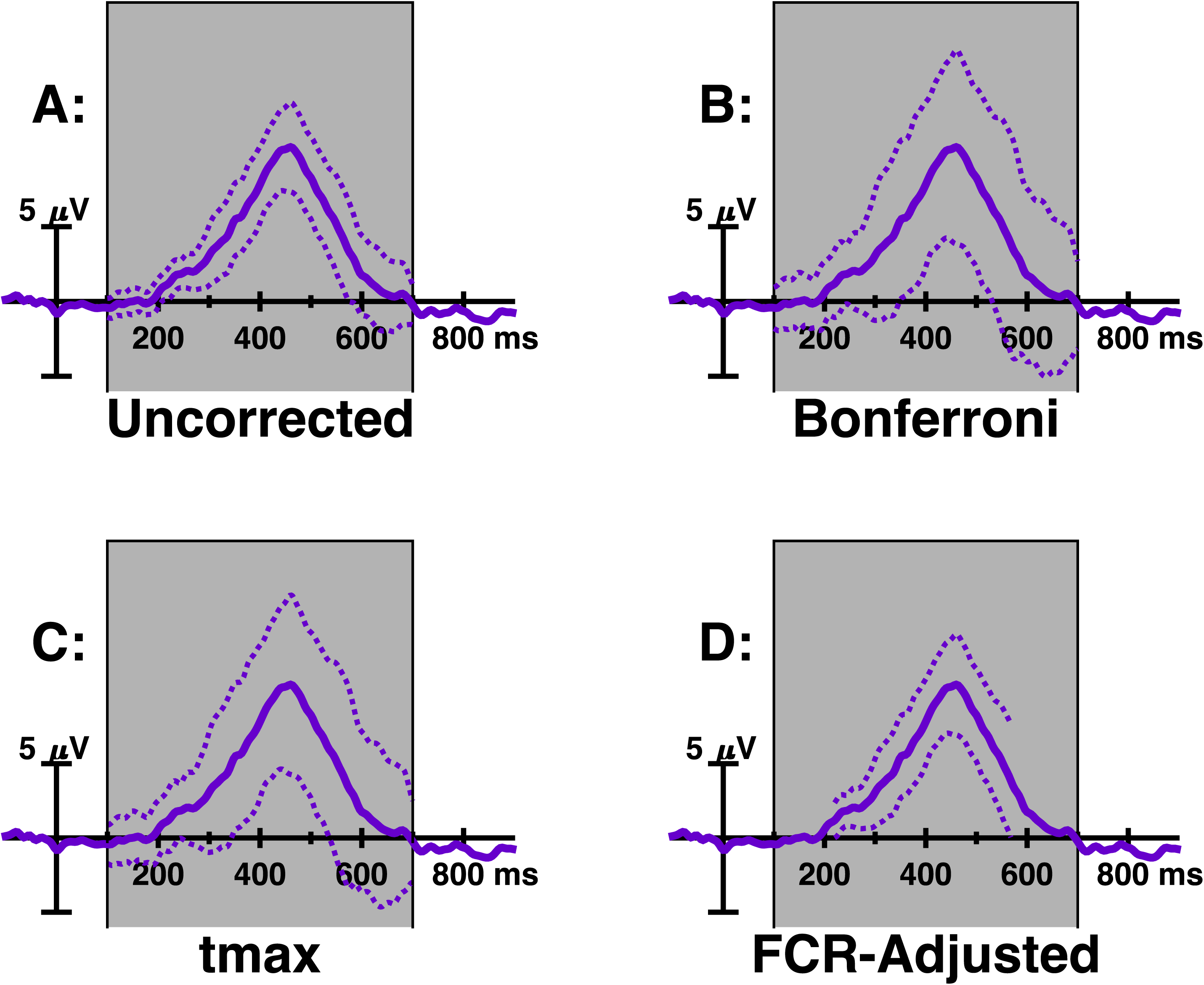
Four different ways of deriving 95% CIs for the target-standard difference wave introduced in Figure 2:B. Solid line represents the difference wave ERP and dashed lines represent 95% confidence intervals. Grey box indicates the time window of interest in which statistical inference is done. The *tmax* resampled CIs were derived using the permutation algorithm described in the text. FCR-adjusted CIs were derived after BH-selection. Note that for the FCR-adjusted BH-selected CIs, we are not able derive CIs for all time points of interest.

### tmax Resampled Confidence Intervals

Although Bonferroni correction is simple to understand and implement, it is too conservative if the multiple parameters being estimated are not independent. Since EEG data are typically highly correlated across nearby time points and scalp locations, Bonferroni correction of EEG analyses is usually extremely over-conservative. For such data, it is generally better to use resampling methods to correct for multiple comparisons. These methods use permutations of the data or bootstrap samples (i.e., sampling from the data with replacement) to estimate the likelihood of getting extreme values, which can then be used to derive *p*-values and CIs. Because the resampled data reflects the degree of dependency between the multiple variables, these methods can achieve exactly the desired level of protection from erroneous inferences and are typically much more powerful than Bonferroni correction (Groppe, Urbach, & Kutas, 2011a; 2011b).

Using *tmax* (Blair & Karniski, 1993) or cluster-based (Bullmore et al., 1999; Maris & Oostenveld, 2007) permutation test are relatively popular among ERP researchers for computing multiple comparison corrected *p*-values (for review see Groppe et al., 2011a). There is not yet a clear analog to cluster-based tests for deriving CIs, but there is for the *tmax* procedure. It is based on bootstrap resampling from the *residuals* of the data, which is the variability in the data that cannot be explained by the statistical model. For example, the procedure for deriving symmetric CIs for estimates of the mean of a family of variables is (Westfall & Young, 1993):

1. For each variable, *y_z_*, remove the estimated mean of each variable, 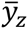, from each data point to derive the residuals, 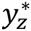. Do this for each variable, *z*, and observation, *x*:

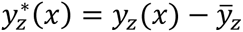
2. Sample *N* times from the residuals with replacement and compute a *t*-score for each variable, where *N* is your sample size. In the equation below, *w* indicates the *w*th such sample, 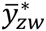 is the mean and 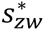 is the standard error of that sample for the *z*th variable.

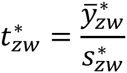
3. From the set of *m t*-scores, where *m* is the number of variables, record the most extreme *t*-score, *tmax_w_*.
4. Repeat Steps 2–3 a large number of times (e.g., 5000) to derive a bootstrap distribution of *tmax*.
5. Find the 100 * (1 *− α*) percentile of the absolute value of the *tmax* distribution, *tmax*_1−_*_α_*.
6. Derive your *tmax*_1−_*_α_* CIs for each variable using that value: 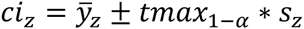

This method is *asymptotically accurate*, in that it will become increasingly accurate as the sample size increases but may be inaccurate for small sample sizes. Simulated ERP data (see Supplemental Materials) suggest that this method is much too conservative for sample sizes of 23 participants or less. A more accurate method (see Supplemental Materials) is to use permutations of the signs of the residuals rather than bootstrap samples. Specifically Step 2 and 4 in the above algorithm are replaced by:

2. Randomly flip the sign (i.e., positive or negative) of each residual with 50% probability and compute a *t*-score for each variable. In the equation below, *w* indicates the *w*th such permuted-sign sample, 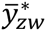 is the mean and 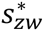 is the standard error of that sample for the *z*th variable.

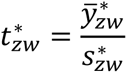
4. Repeat Steps 2–3 for all *2^N^* possible sets of signs for small sample sizes (e.g., *N*<13) where *N* is your sample size. For larger sample sizes, simply repeat Steps 2–3 a large number of times (e.g., 5000). These repetitions produce a permutation distribution of *tmax*.

Applying this method to simulated ERP data (see Supplemental Materials) to generate 95% CIs, suggests that it provides accurate control of the familywise error rate for samples consisting of as few as 9 participants. For extremely small sample sizes the small number of possible permutations limits the CI coverages that can be accurately estimated. For example, with four participants, there are only 16 possible permutations and all percentiles above 87.5% (i.e., 1-2/16) are equivalent.

To illustrate, we apply this method to the visual oddball difference wave from 100 to 700 ms and get a t*max*_0.95_ value of 4.25. Figure 3:C illustrates the resultant CIs, which are smaller than those derived with the Bonferroni-method but provide the same degree of protection against false CI coverage as Bonferroni CIs. Note that with more comparisons (e.g., tests at every time point from 100 to 700 ms at multiple electrodes) the difference between Bonferroni and *tmax* CIs should increase.

### False Discovery Rate-Corrected Confidence Intervals

Another, more powerful and somewhat more permissive alternative to the *tmax* procedure for multiple comparison correction is to control the false discovery rate (FDR) of the family of tests. While Bonferroni and *tmax* correction control the probability of obtaining one or more false positive test results, FDR control limits the proportion of positive test results that are false positives. This more permissive criterion makes FDR control generally quite powerful while still providing weak control of the familywise error rate. This means that if any effects are detected, you can be as sure as if you had done Bonferroni correction that some effect is truly present but there is a higher chance that individual comparisons are false positives.

There are several algorithms for FDR control, but the most popular are that of Benjamini and Hochberg (1995) and Benjamini and Yekutieli (2001). The Benjamini and Hochberg (BH) procedure operates as follows:

1. Sort the *p*-values from the entire family of *m* tests (i.e., *m* is the total number of hypothesis tests) from smallest to largest (*p_i_* refers to the *i*th smallest *p*-value).
2. Define *k*, as the largest value of *i* for which the following is true:

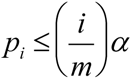
3. If at least one value of *i* satisfies this relationship, then hypotheses 1 though *k* are rejected, otherwise no hypotheses are rejected.

This procedure guarantees that FDR will be less than or equal to α, if the different tests being compared are independent or exhibit positive regression dependency. For Gaussian distributed data (as ERPs likely approximately are), positive regression dependency means that all the variables being tested are positively correlated or uncorrelated. In practice, however, the BH algorithm may effectively control FDR even if some variables are negatively correlated. Theoretical results show that when data come from relatively light-tailed distributions (e.g., Gaussian), FDR control performs as if the tests were independent as the number of tests in the family increases (Clarke & Hall, 2009). Thus with a sufficient number of tests, FDR procedures guaranteed to work when the tests are independent should also provide accurate control of FDR for ERP data. In fact, applications of the BH FDR procedure to simulated ERP data sets have found that it does control FDR at or below the nominal level despite some variables being negatively correlated (Groppe et al., 2011b).

The more conservative Benjamini and Yekutieli (BY) algorithm is the same as BH but Step 2 is replaced by:

2. Define *k*, as the largest value of *i* for which the following is true:

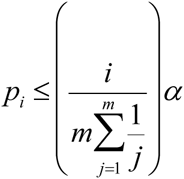

The BY algorithm is always guaranteed to control the FDR, but may be overly conservative in practice.

These FDR control algorithms can also be used to adjust CIs for multiple comparisons as follows (Benjamini & Yekutieli, 2005):

1. Apply the BH or BY FDR procedure to the *p*-values from your family of tests.
2. For any *p*-values that are significant after FDR correction, construct a CI for the corresponding test with coverage 1-α’, where α’ is:

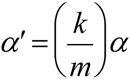

These CIs are called false coverage-statement rate-adjusted BH- or BY-selected CIs. False coverage-statement rate (FCR) is the proportion of constructed CIs that do not cover the true value of the parameter. FCR-adjusted BH-selected CIs guarantee the expected FCR is less than or equal to α if the different tests being compared are independent or exhibit positive regression dependency. FCR-adjusted BY-adjusted selected CIs always guarantee that the expected FCR is less than or equal to α. Moreover, there is a intuitive duality between the FCR-adjusted selected CIs and FDR adjusted *p*-values: any tests with significant FDR adjusted *p*-values will have FCR-adjusted selected CIs that do not include the null hypothesis value.

Figure 3:D illustrates the FCR-adjusted BH-selected CIs for the visual oddball data. In this case the difference wave at 88 of the 151 time points of interest significantly differ from 0 after FDR-BH correction. This means that our FCR-adjusted BH-selected CI coverage needs to be 97.1%. These CIs are much smaller than those derived with the Bonferroni or *tmax* adjustment. However, we don’t have CIs for all time points in our window of interest because the BH-adjusted *p*-values at those time points were not less than 0.05. This can be a serious limitation of FCR-adjusted CIs. Non-significant *p*-values could indicate that there truly is no effect of consequence at that variable or they could indicate that there is insufficient statistical power to detect the effect. FCR-adjusted CIs cannot distinguish between these alternatives. An additional shortcoming of FCR-adjusted CIs is that we can be less certain of the accuracy of any single CI than if we had used Bonferroni or *tmax* adjustment. However, on average, if we repeated this experiment and analysis many times, the great majority of the CI’s would be accurate using conventional *p*-value thresholds.

## C. Summary

Statistical inference in psychophysiological studies have conventionally focused on null hypothesis testing (i.e., *p*-values) and often not reported confidence intervals (CIs). CIs provide upper and lower bounds that span the true effect being studied with a known probability. This information is complementary to that communicated by *p*-values. CIs help us to see if a statistically significant result might have been generated by a small, inconsequential effect or if a sizable effect might have been missed by insufficient statistical power. Moreover, CIs can help plan future studies by providing a sense of how many observations will be needed to detect that effect again and can facilitate comparing different studies. In this article, I have illustrated how to derive CIs in ways that compensate for multiple comparisons using an event-related potential (ERP) time series. CIs can also be derived via these methods for ERP topographies (e.g., Figures 4–5) and any other psychophysiological measure. Which method to use for computing CIs depends on the goals of the analysis and the required degree of certainty. Recommendations for when to use the four methods covered in this review are summarized Table 1.

**Figure 4:**
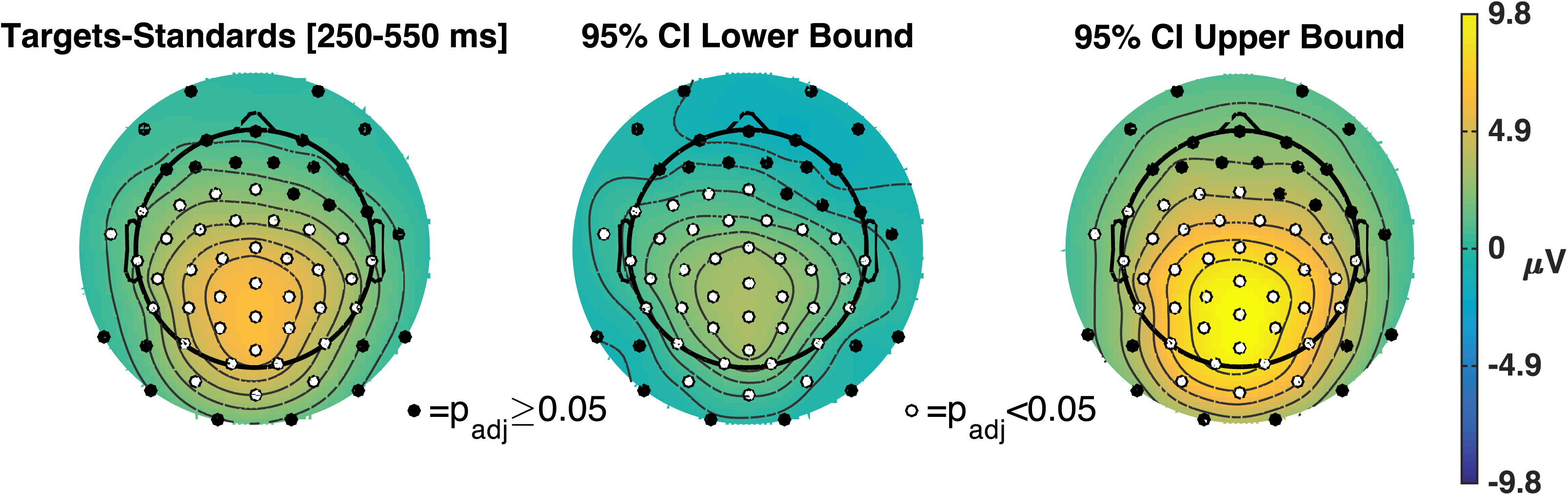
[Left] Mean targets-standards difference wave across 63 EEG channels from 250 to 550 ms post-stimulus onset. [Middle-Right] max-derived 95% CIs on the topography of the mean difference wave.

**Figure 5:**
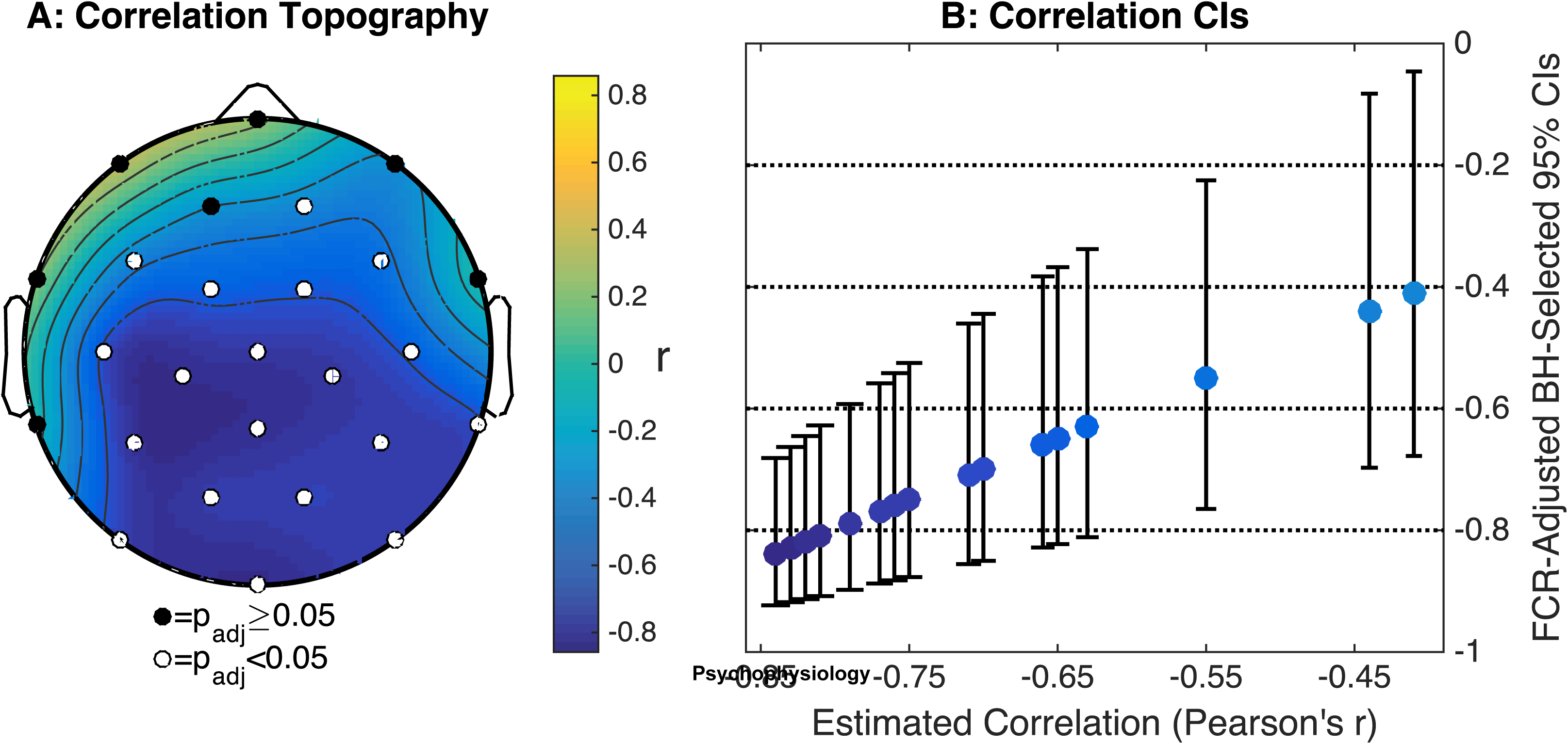
An example of how to report CIs for topographies of statistics like Pearson’s correlation coefficient, for which the Ci is simply a function of the statistic. Such plots are called “confidence calibration plots” (Rosenblatt & Benjamini, 2014).[Left] Topography of linear correlations between mean ERP amplitude from 200–500 ms following word onset and the cloze probability of that word (data from (DeLong, Urbach, & Kutas, 2005). Electrode colors indicate statistical significance after Benjamini and Hochberg FDR correction for the 26 statistical tests. [Right] FCR-adjusted BH-selected 95% CIs of the ERP x cloze probability correlation for the 19 electrodes with significant correlations.

**Table 1:** Situations in which each of the methods for deriving confidence intervals reviewed here are best suited.

| Confidence Interval Type | Recommended Conditions of Use |
| --- | --- |
| No Correction for Multiple Comparisons | • Intervals are derived for weakly exploratory purposes, to simply provide a sense of data variability, or to illustrate the strength of a null hypothesis (i.e., CIs clearly span the null hypothesis even when not correcting for multiple comparisons) |
| Bonferroni-Corrected CIs | • Accurate bounds are needed on every single variable (e.g., for clearly demonstrating the latency of effect onset)<br>• Sample size is too small for resampled CIs |
| *t*<sub>max</sub> Resampled CIs | • Useful in the same situations as Bonferroni-corrected CIs and sample size is sufficiently large (e.g., nine or more participants for permutation method) |
| FCR-Adjusted BH-Selected CIs | • Goal is to get a general sense of when and where effects occur, but precise effect boundaries are not needed (i.e., accurate bounds are *not* needed on every single variable)<br>• Some effects are expected and CIs are not needed where there is not significant evidence of an effect |
| FCR-Adjusted BY-Selected CIs | • Useful in the same situations as FCR-adjusted BH-selected CIs but provides a greater degree of certainty as to CI accuracy (i.e., accuracy is guaranteed and a smaller number of CIs will typically be incorrect) |

If CIs become a conventional part of psychophysiological research, it will reduce the scientific decline effect by helping us to better understand just how reliable our results are to begin with and to focus on finding more effects that stand the test of time. Of course, CIs alone will not fix the scientific decline effect. Selective reporting of results (i.e., the file drawer effect) and post-hoc changes in data acquisition and analysis will still lead to overestimation of effect sizes even if CIs (and other inferential statistical tools) are used with the best of intentions. Moreover, these biases will continue to be severe as long as underpowered studies remain the norm in the neurosciences.

To facilitate the use of CIs in ERP studies, I have provided Matlab code for deriving and visualizing CIs for ERPs using the approaches described here as part of the Mass Univariate ERP Toolbox (http://openwetware.org/wiki/Mass_Univariate_ERP_Toolbox). This code covers the common cases of computing CIs for a single set of ERPs or difference waves. The *multcomp* package for *R* (https://cran.r-project.org/web/packages/multcomp/index.html) also implements these methods for computing CIs and can handle more complicated analyses (e.g, multiple linear regression).

**Author Notes:** The author has no conflicts of interest regarding this manuscript and would like to thank Dr. Suzanne Wood, Dr. Peter Westfall, Dr. Michael J. Larson, and an anoynmous reviewer for assistance with this work (though their assistance should not be interpreted as endorsements). Please contact the author at for reprints.

## Supplemental Materials: *tmax* Resampled CIs of Simulated ERP Data

The accuracy of the *tmax* resampling procedure requires having a sample size that is large enough to sufficiently approximate the population being sampled from. To get a sense of what “large enough” is for ERP data, I applied the procedure to simulated ERP data and quantified its accuracy as a function of the number of participants.

### Simulation Methods

ERP data were simulated using a procedure and data set used previously (Groppe, Urbach, & Kutas, 2011). Specifically, realistic EEG background noise was derived from the data of 23 volunteers who performed a linguistic priming task (Groppe, Choi, Topkins, & Kutas, 2009). The University of California, San Diego Institutional Review Board approved the experimental protocol. Each participant’s EEG was recorded at 26 scalp channels using a left mastoid reference and an analog bandpass filter of 0.016–100 Hz. EEG was digitized at a 250 Hz sampling rate. After recording, the EEG was re-referenced to the algebraic mean of both mastoids, low-pass filtered at 50 Hz, and artifact polluted trials were either rejected or artifact corrected using independent components analysis (Lee, Girolami, & Sejnowski, 1999). ERPs were derived from epochs of EEG time-locked to tones and lasting from −100 to 920 ms peri-tone onset. The ERP was then subtracted from each epoch to produce trials of zero mean EEG background noise. On average, there were 223 trials per participant (SD=12). Variables of interest for these simulations were all time points from 100 to 900 ms at all 26 scalp channels for a total of 5226 dependent variables (i.e., 26 channels × 201 time points). The median standard deviation of the background noise at these data points across all 23 participants was 10.56 μV (IQR=3.41 μV).

To simulate a single ERP experiment, data from 23 participants were randomly selected without replacement. ERPs were derived for each of the participants by randomly selecting (with replacement) 49 of that participant’s background noise trials, removing the mean prestimulus voltage (−100 to 0 ms), and averaging the trials. A deflection of 1.3 μV at 13 central and posterior electrodes was added from 400 to 700 ms to simulate a “P300-like” effect (Bentin, Mouchetant-Rostaing, Giard, Echallier, & Pernier, 1999). This procedure was repeated 5000 times for four different sample sizes: 4, 9, 16, and 23 participants. CIs were generated for each variable of interest using the two *tmax* resampling methods described in this manuscript: [1] bootstrap resampling of residuals and [2] permutations of residual signs. For the boostrap procedure, 5000 boostrap samples were used. For the permutation procedure all possible permutations were used for sample sizes smaller than 16 participants. For 16 or more participants, 5000 random permutations were used.

Data and Matlab code for performing these simulations are publicly available via the Open Science Framework (https://osf.io/5tmwp/).

### Simulation Results

For each sample size and method, I computed the familywise error rate (FWER) of the CIs (i.e., the proportion of simulations for which one or more CI did not span the true ERP mean). Results are shown in Supplemental Figure 1. For these sample sizes, all methods control FWER below the nominal 5% rate. Bonferroni FWER is generally far below 5% because it does not account for the fact that the data at nearby electrodes and time points are highly correlated. The *tmax* bootstrapped residual CIs are even more conservative than the Bonferonni CIs for these sample sizes, but would become increasingly accurate with more participants. In contrast, the *tmax* permuted residual CIs accurately control FWER for all but the smallest sample size. For such small sample sizes, the limited number of permutations (in this case 16) necessarily makes the method overly conservative.

**Supplemental Figure 1:**
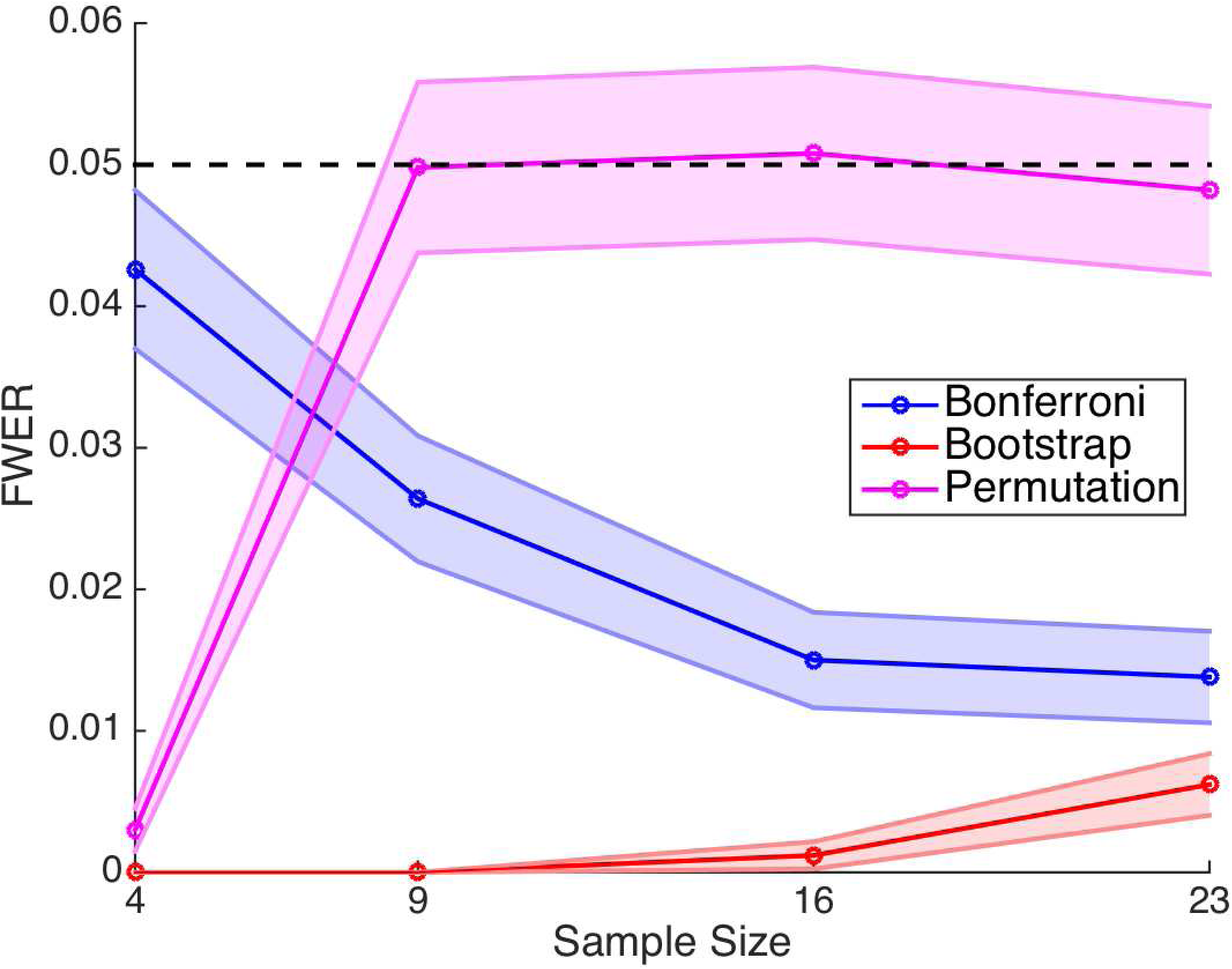
Familywise error rate (FWER) of 95% CIs derived from 5000 simulated ERP datasets as a function of sample size (i.e., number of participants). Three methods for correcting CIs for multiple comparisons were used: Bonferroni, *tmax* bootstrapped residuals, *tmax* permuted residuals. Dashed line indicates the nominal 5% error rate. Shading indicate 95% uncorrected CIs.

